# mTORC2/AKT/HSF1/HuR constitute a feed-forward loop regulating Rictor expression and tumor growth in glioblastoma

**DOI:** 10.1101/140293

**Authors:** Brent Holmes, Angelica Benavides-Serrato, Ryan S. Freeman, Kenna A. Landon, Tariq Bashir, Robert N. Nishimura, Joseph Gera

## Abstract

Overexpression of Rictor has been demonstrated to result in increased mTORC2 nucleation and activity leading to tumor growth and increased invasive characteristics in glioblastoma multiforme (GBM). However the mechanisms regulating Rictor expression in these tumors is not clearly understood. In this report, we demonstrate that Rictor is regulated at the level of mRNA translation via HSF1-induced HuR activity. HuR is shown to directly bind the 3′ UTR of the Rictor transcript and enhance translational efficiency. Moreover, we demonstrate that mTORC2/AKT signaling activates HSF1 resulting in a feed-forward cascade in which continued mTORC2 activity is able to drive Rictor expression. RNAi-mediated blockade of AKT, HSF1 or HuR is sufficient to downregulate Rictor and inhibit GBM growth and invasive characteristics *in vitro* and suppresses xenograft growth in mice. We further demonstrate that constitutive overexpression of HuR is able to maintain Rictor expression under conditions of AKT or HSF1 loss. In an additional level of regulation, *miR-218*, a known Rictor targeting miRNA is shown to be subject to mTORC2/STAT3-mediated repression. The expression of these components is also examined in patient GBM samples and correlative associations between the relative expression of these factors support the presence of these signaling relationships in GBM. These data support a role for a feed-forward loop mechanism by which mTORC2 activity stimulates Rictor translational efficiency and suppresses *miR-218* resulting in enhanced mTORC2 activity in these tumors.

## INTRODUCTION

The mechanistic target of rapamycin (mTOR) protein kinase integrates signal transduction networks coordinating cell growth, nutrient status, protein synthesis and autophagy ^1^. mTOR exists in two functionally distinct complexes, mTORC1 and mTORC2 ^2^. The mTORC1 and mTORC2 kinase complexes both share mTOR, mLST8 (GβL), Deptor and Tti1/Tel2, however mTORC1 also specifically contains Raptor and PRAS40, while mTORC2 contains the subunits Rictor, mSIN1 and Protor1/2 ^3^. The regulatory signals impinging on mTORC1 activity have been the focus of many studies, however by contrast, the mechanisms of mTORC2 regulation are not well understood. mTORC2 is activated by growth factor receptors and PI3K signaling ^4^ Association with the ribosome itself has additionally been demonstrated to result in mTORC2 activation ^5, 6^. mTORC2 has been shown to activate several downstream substrates, however its most well characterized function is the phosphorylation of serine 473 of AKT within the hydrophobic turn motif resulting in full activation of AKT ^7^. In GBM, the mutated epidermal growth factor receptor (EGFR) variant, EGFRvIII is known to activate mTORC2, in addition to PTEN loss ^8-10^. In an EGFR-PI3K driven *Drosophila* glial tumor model mTORC2 activity was required for GBM formation, and Rictor overexpression in a GEMM was sufficient to induce gliomagenesis ^11, 12^.

Rictor is a 200 kD protein initially identified as the defining component of mTORC2 and lacks any significant sequence homology between yeast and mammals ^4, 13^. Moreover, Rictor lacks any structural domains of known function but contains a C-terminus which is conserved among vertebrates. The degree of Rictor association with mTOR appears to be inversely correlated to Raptor expression and varies in different cell types ^7, 13^. Rictor is overexpressed in several cancers leading to hyperactive mTORC2 and has been shown to play a causal role in glioma formation ^12, 14-17^. Rictor expression has been demonstrated to be regulated transcriptionally and via protein degradation ^18, 19^, however recent studies have suggested that Rictor expression may also be regulated post-transcriptionally ^20^.

Here we describe a feed-forward cascade involving activation of AKT/HSF1 resulting in HSF1-induced HuR expression. Rictor is demonstrated to be a target of HuR leading to enhancement of Rictor mRNA translation and elevated mTORC2 activity. Rictor mRNA is demonstrated to be subject to translational control and shown that HuR binds to the Rictor 3′ UTR enhancing translational efficiency. Data is shown which demonstrate that HuR is a direct target of HSF1. Knockdown of AKT, HSF1 or HuR results in down-regulation of Rictor expression and impedes GBM growth, migration and invasive properties. Uncoupling HuR expression from its native promoter via viral expression maintained Rictor expression under conditions of AKT or HSF1 loss. Furthermore, data is shown supporting the repression of *miR-218* expression by elevated mTORC2 via STAT3 and examination of clinical GBM specimens support the proposed signaling relationships.

## RESULTS

### Rictor is regulated at the level of mRNA translation

To begin investigating the mechanism(s) of Rictor expression in GBM we examined steady-state mRNA and protein levels in U138 GBM cells following stimulation. This line expresses relatively low levels of Rictor and we reasoned that signaling which induces expression would be readily discernable. We treated cells with epidermal growth factor (EGF) and monitored mRNA and protein levels at time points following stimulation. As shown in figure 1A (*left panel*), EGF treatment did not significantly alter the steady-state levels of Rictor mRNA, suggesting enhanced transcription or effects on mRNA stability did not significantly regulate Rictor expression, however we did observe a marked increase in protein levels (Fig. 1A *middle and right panels*). To examine the possibility that post-translational degradation of Rictor was inhibited by EGF exposure, we monitored the turnover of Rictor protein in the absence and following exposure to EGF. As shown in figure 1B, EGF treatment did not alter the degradation of Rictor. We also performed polysome analysis on the Rictor mRNA following EGF stimulation and as shown in figure 1C, Rictor mRNA displayed a marked shift to polysome-containing sucrose density gradient fractions consistent with an increase in the translational efficiency of this transcript and the accumulation of Rictor protein following EGF exposure. These data demonstrate that the Rictor mRNA is subject to translational regulation following EGF treatment.

**Figure 1.**
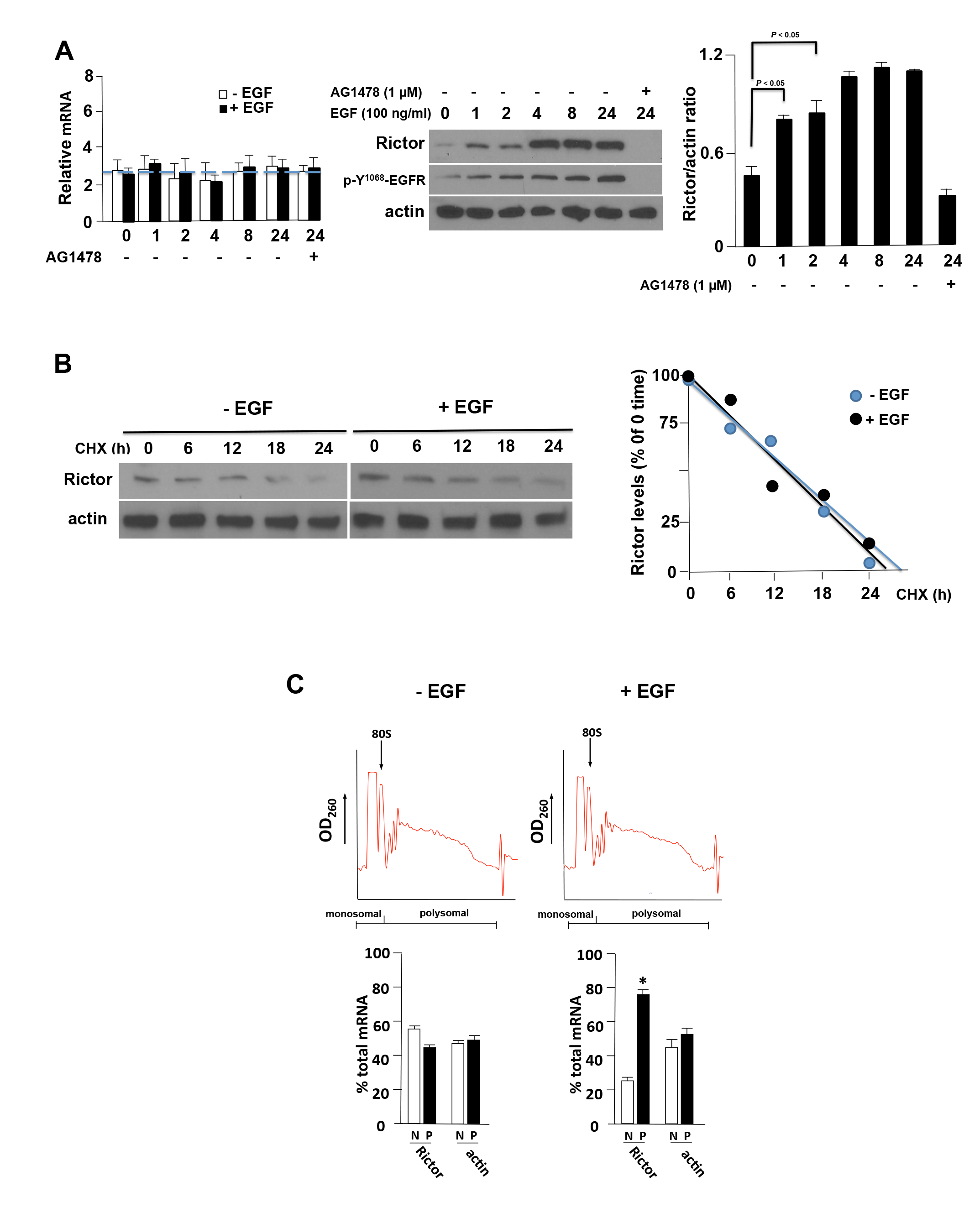
Rictor mRNA is regulated at the level of translation. (**A**) Steady-state Rictor mRNA levels in U138 cells in the absence or presence of EGF (100 ng/ml) for the indicated time points (h) (*left panel*). U138 cells were treated as indicated with EGF or in combination with the EGFR inhibitor AG1478 (5 μM; negative control). Mean + S.D. are shown, n = 3. Rictor protein accumulation in U138 stimulated with EGF or AG1478 as indicated (*middle panel*). Cell lysates were subjected to immunoblot analyses for the indicated proteins and blots were quantified by densitometry and results depicted in the *right panel*. Mean + S.D. are shown, n = 3. (**B**) Rictor protein stability is not affected by EGF stimulation. U138 cells were subjected to cycloheximide (CHX)-chase experiments and Rictor protein levels examined at the indicated time points in the absence or presence of EGF (100 ng/ml) (*left panel*). Band intensities were quantified by densitometry and displayed graphically (*right panel*). (**C**) Polysome distribution of Rictor and actin mRNAs. U138 cells were treated with EGF (100 ng/ml, 8 h) and extracts subjected to sucrose density gradient centrifugation and divided into 11 1-ml fractions which were pooled into nonribosomal, monosomal fraction (N, white bars) and a polysomal fraction (P, black bars). Purified RNAs were used in real time quantitative rt-PCR analyses to determine the distributions of Rictor and actin mRNAs across the gradients. Polysome tracings are shown above values obtained from the rt-PCR analyses which are displayed graphically below. rt-PCR measurements were performed in quadruplicate and the mean and + S.D. are shown.

### HuR binds to the 3′ UTR of Rictor mRNA and promotes translation

To understand the mechanism by which Rictor mRNA translation is regulated we initially examined the structure of the Rictor transcript. In a computational survey of RNA-binding protein motifs, we identified four consensus HuR binding sites located within the 3′ UTR of the Rictor mRNA which were conserved in human and mouse transcripts (Fig. 2A). These sites were found to be significantly more U-rich than AU-rich and adhered to the consensus sequences derived from studies by López de Silanes *et al* ^21^ and separately by Tenenbaum and coworkers ^22^. As HuR is known to enhance mRNA stability and translation ^23^ we first determined whether HuR would bind to these sequences. In an RNA pull-down assay, extracts from U138 cells treated without or with EGF, were mixed with biotinylated HuR binding site motif(s) RNA sequences as shown in Figure 2B. HuR was preferentially precipitated by each of the HuR binding site motifs (1-4) in a manner dependent on EGF stimulation. HuR was not detected in samples which were precipitated by a nonspecific control RNA. Similarly, in HuR immunoprecipitates we were able to detect Rictor 3′ UTR sequences by rt-PCR, which was enhanced in extracts treated with EGF (Figure 2C). To investigate whether these Rictor 3′ UTR HuR binding motifs were involved in regulating translational efficiency, we generated heterologous reporter mRNA transcripts in which the full-length Rictor 3′ UTR was fused to the luciferase ORF and the HuR binding motifs mutated (Figure 2D). The effects of these mutations on mRNA translation were subsequently assessed. As shown in figure 2D, the full-length Rictor 3′ UTR containing mRNAs were markedly shifted to polysomal fractions by EGF stimulation consistent with enhanced HuR binding. However, mutating motifs 1 and 2 reduced the amount of reporter mRNA which was polysome associated and mutating all four HuR binding motifs completely abolished EGF-stimulated mRNA reporter polysome association. Taken together these data demonstrate that the mRNA translational enhancer HuR binds to the Rictor 3′ UTR to stimulate translational efficiency.

**Figure 2.**
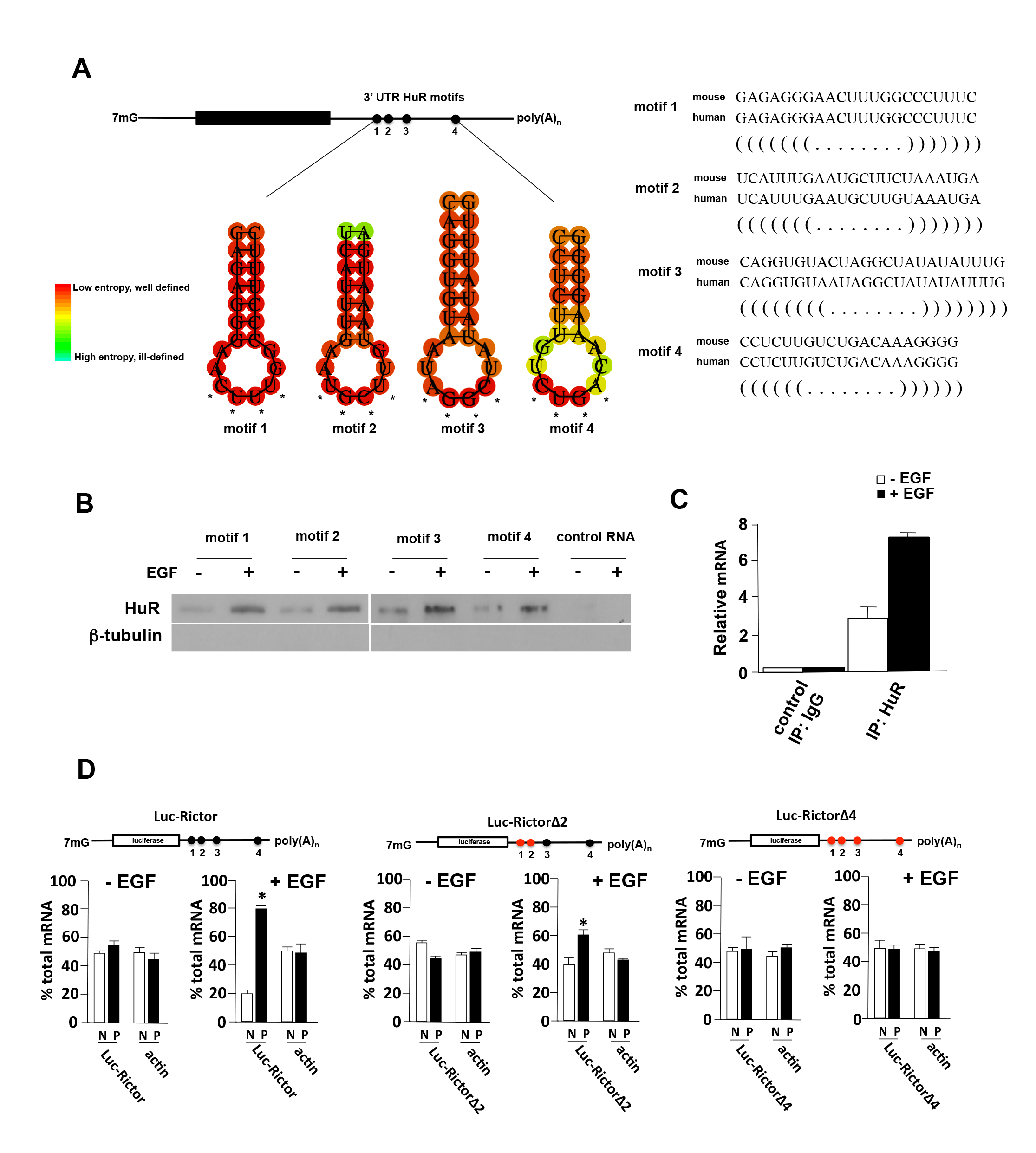
HuR binds to the 3′ UTR of Rictor and stimulates translation. (**A**) Sequence and structure of predicted HuR binding motifs within the Rictor 3′ UTR (*left panel*). Structural alignment of mouse and human HuR motifs within the Rictor 3′ UTRs (*right panel*). (**B**) Identification of HuR in RNA pull-down assays utilizing biotinylated HuR binding motifs as indicated. Biotinylated nonspecific RNA was used as a negative control. Cytoplasmic extracts of U138 cells treated in the absence or presence of EGF (100 ng/ml, 8 h) were incubated with biotinylated RNAs corresponding to HuR binding motifs 1-4 as shown and precipitated with streptavidin-Sepharose beads. Bound fractions were analyzed by immunoblotting for HuR and tubulin. Experiments were repeated three times with similar results. (**C**) HuR binds to Rictor 3′ UTR RNA containing HuR binding motifs 1-4 in cells and binding is stimulated by EGF (100 ng/ml, 8 h). Control IgG or anti-HuR antibody was used to immunoprecipitate (IP) lysates from U138 cells, and bound RNA was amplified by PCR of the Rictor 3′ UTR sequences. Relative amounts of Rictor 3′ UTR RNA are displayed graphically. Mean + S.D. are shown, n = 3. (**D**) Polysome analyses of Luc-Rictor 3′ UTR containing reporter mRNAs. U138 cells expressing the indicated reporter mRNAs, native Rictor 3′ UTR (Luc-Rictor), mutant 1-2 (Luc-RictorΔ2) in which HuR binding motifs 1 and 2 were mutated, and mutant 1-4 (Luc-RictorΔ4) in which all four HuR binding motifs were mutated (4 unpaired bases were changed to C within the loop structures of the predicted hairpin loop motifs, see Fig. 2A, asterisks), were treated with EGF (100 ng/ml, 8 h) and lysates subjected to polysome analyses as in Fig. 1C. Purified RNAs were used in real time quantitative rt-PCR analyses to determine the distributions of Luc-Rictor and actin mRNAs across the gradients. Values obtained from the rt-PCR analyses are displayed graphically below. rt-PCR measurements were performed in quadruplicate and the mean and + S.D. are shown.

### mTORC2/AKT/HSF1 signaling stimulates HuR transcription in GBM

Overexpression of Rictor in glioblastoma cells has been demonstrated to increase the nucleation of mTORC2 resulting in elevated kinase activity ^12, 16^. We sought to identify a possible signaling mechanism linking the mTORC2/AKT axis to the regulation of HuR. AKT has been shown to directly activate HSF1 via phosphorylation of serine 326 ^24^ Additionally, Chou *et al* have recently demonstrated that HSF1 appears to regulate *β*-catenin expression via effects on HuR in breast cancer cells ^25^. These relationships suggested to us that mTORC2/AKT signaling may activate HSF1/HuR leading to enhanced translation of Rictor which in turn, would enhance mTORC2 activity in a feed-forward loop promoting GBM proliferation and motility. We examined these signaling cascades in three GBM cell line models of elevated mTORC2 activity. mTORC2 is known to be activated by EGF stimulation or overexpression of the mutated constitutively active EGFRvIII allele, as well as, by overexpression of Rictor ^8, 12, 16, 26^. As shown in Figure 3A, in U138 cells stimulated with EGF, phospho-S^473^-AKT, phospho-S^326^-HSF1, HuR and Rictor levels were enhanced following stimulation (see also Supplemental Figure S1). In H4 cells in which Rictor is overexpressed ^16^, phospho-S^473^-AKT, phospho-S^326^-HSF1 and HuR expression was enhanced relative to the levels of these proteins in parental H4 cells. Similarly, in U87 cells overexpressing the mutant EGFRvIII allele, phospho-S^473^-AKT, phospho-S^326^-HSF1, HuR and Rictor protein levels were elevated compared to parental U87 cells. While these data supported the notion of an operative mTORC2/phospho-S^473^-AKT/phospho-S^326^-HSF1/HuR/Rictor feed-forward signaling cascade, the mechanism by which activated HSF1 leads to increases in HuR expression was unclear in GBM cells. To examine whether HuR was a direct transcriptional target of HSF1 we searched the HuR promoter for canonical or noncanonical heat shock elements (HSEs) ^27^. We identified tandem noncanonical HSEs beginning at position −475 within the HuR promoter and these sequences were conserved in the mouse HuR promoter (Figure 3B). Interestingly, Mendilo *et al* also identified this region of the HuR promoter as a potential binding target of HSF1 in a ChIP-Seq study of breast cancer cells ^28^. To determine whether these candidate HSEs were capable of mediating transcriptional responses directed by HSF1, we determined both RNA *Pol* II and HSF1 occupancy via ChIP assays followed by quantitative PCR. As shown in Figure 3C (*left panel*), *Pol* II content within the HuR promoter containing the tandem HSEs was increased in U138 cells following EGF stimulation. There was also a marked enhancement of *Pol* II content within the HuR promoter in H4_Rictor_ - or U87EGFRvIII-overexpressing cells relative to parental cells. Similar results were obtained analyzing HSF1 occupancy of the HuR promoter as EGF stimulation, Rictor- or EGFRvIII-overexpression all resulted in marked increases in HSF1 occupancy (Figure 3C, *right panel*). The phosphorylation state of functionally bound HSF1 was also assessed in a series of *in vitro* DNA-pull down assays (Figure 3D). Consistent with the chromatin immunoprecipitation experiments, EGF stimulation, Rictor- or EGFRvIII-overexpression resulted in higher levels of bound phospho-S^326^-HSF1 and total HSF1 compared to unstimulated U138 cells or parental H4 and U87 cells.

**Figure 3.**
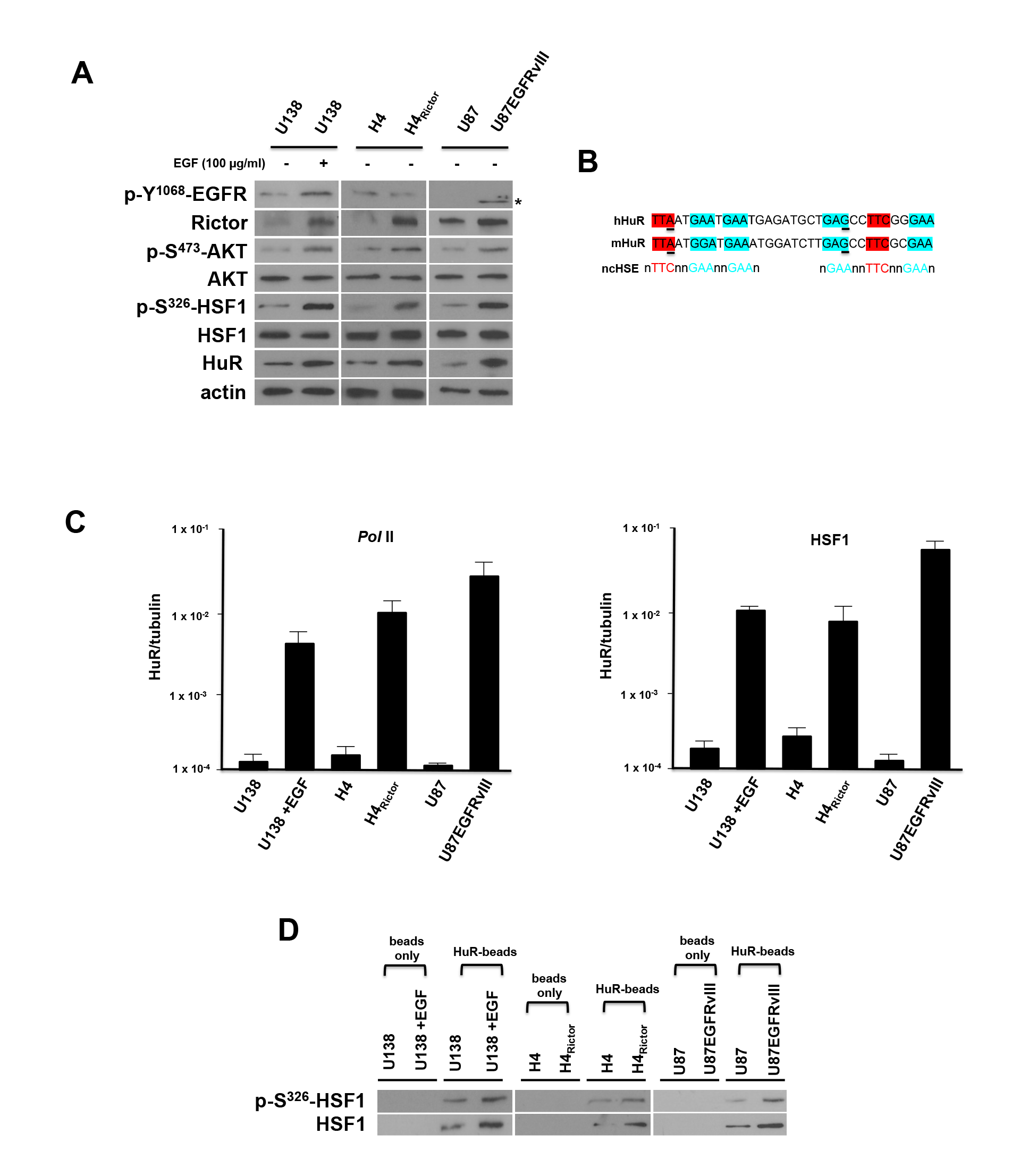
mTORC2 activation leads to HSF1-stimulated HuR transcription. (**A**) Signaling effects of mTORC2 stimulation by EGF, Rictor or EGFRvIII overexpression. The GBM lines were treated with EGF (I00 ng/ml, 8 h) as shown and lysates immunoblotted for the indicated proteins. Asterisk corresponds to the truncated phosphorylated Y^1068^ mutant EGFRvIII. (**B**) Sequence and alignment of tandem noncanonical heat-shock elements (HSEs) identified within the promoter region of HuR (−475 to −441). Previously identified noncanonical HSE (ncHSE) are shown below the human and mouse sequences. Underlined nucleotides differ from those noncanonical HSEs which have been described ^28^. (**C**) ChIP analyses of HuR promoter activity and HSF1 association in the indicated GBM cell lines. RNA polymerase II (*Pol* II) association with HuR promoter (*left panel*) and HSF1 association with HuR promoter (*right panel*) were determined and ChIP-quantitative PCR data are expressed as a ratio of HuR to tubulin. Mean and + S.D. are shown, n = 3. (**D**) Sepharose beads were conjugated with the ncHSEs HuR DNA (HuR-beads) or beads without linked DNA and incubated with nuclear extracts from the indicated cell lines. Following recovery by centrifugation and washing of the beads, bound material was analyzed by immunoblot for phospho-S^326^-HSF1 and total-HSF1. These experiments were performed twice with similar results.

### AKT, HSF1 and HuR are required for Rictor expression and their loss inhibits GBM cell line growth, motility and invasiveness

We next investigated whether loss of AKT, HSF1 or HuR altered Rictor and mTORC2 activity. DBTRG-05MG cells, which harbor elevated mTORC2 activity (B. Holmes and J. Gera, unpublished results), were stably transduced with lentiviral vectors expressing shRNAs targeting AKT, HSF1, HuR, Rictor or a control scrambled sequence non-targeting control. Cells expressing the shRNAs were immunoblotted for the signaling constituents predicted in the feed-forward loop. As shown in Figure 4A (see also Supplemental Figure S2), knockdown of AKT resulted in a decline of phospho-S^326^-HSF1, HuR and Rictor consistent with a signaling pathway involving a concerted AKT/phospho-S^326^-HSF1/HuR/Rictor cascade. Similarly, knockdown of HSF1 resulted in down-regulation of HuR, Rictor and phospho-S^473^-AKT levels, while knockdown of HuR led to reduced expression of Rictor, phospho-S^473^-AKT and phospho-S^326^-HSF1. Knockdown of Rictor abrogated mTORC2 activity and resulted in reduced phospho-S^473^-AKT levels, consistent with previous results ^16^, but also reduced phospho-S^326^-HSF1 and HuR expression. We additionally determined the effects of knockdown of each of these signaling components on growth, migration and invasive characteristics in the lines. As shown in Figure 4B, growth of AKT, HSF1, HuR and Rictor shRNA-expressing lines was markedly reduced as compared to control scrambled non-targeting shRNA-expressing cells. The migratory capacity (Figure 4C) was also significantly reduced in these knockdown lines as loss of AKT, HSF1, HuR and Rictor impeded the ability of cells to migrate on either vitronectin or fibronectin in Boyden chambers. The ability of AKT, HSF1, HuR or Rictor knockdown cells to migrate was reduced by ~ 60-70% relative to control scrambled non-targeting shRNA expressing cells. As shown in Figure 4D, AKT, HSF1, HuR and Rictor knockdown cells were also inhibited in their ability to invade Matrigel relative to control cells.

**Figure 4.**
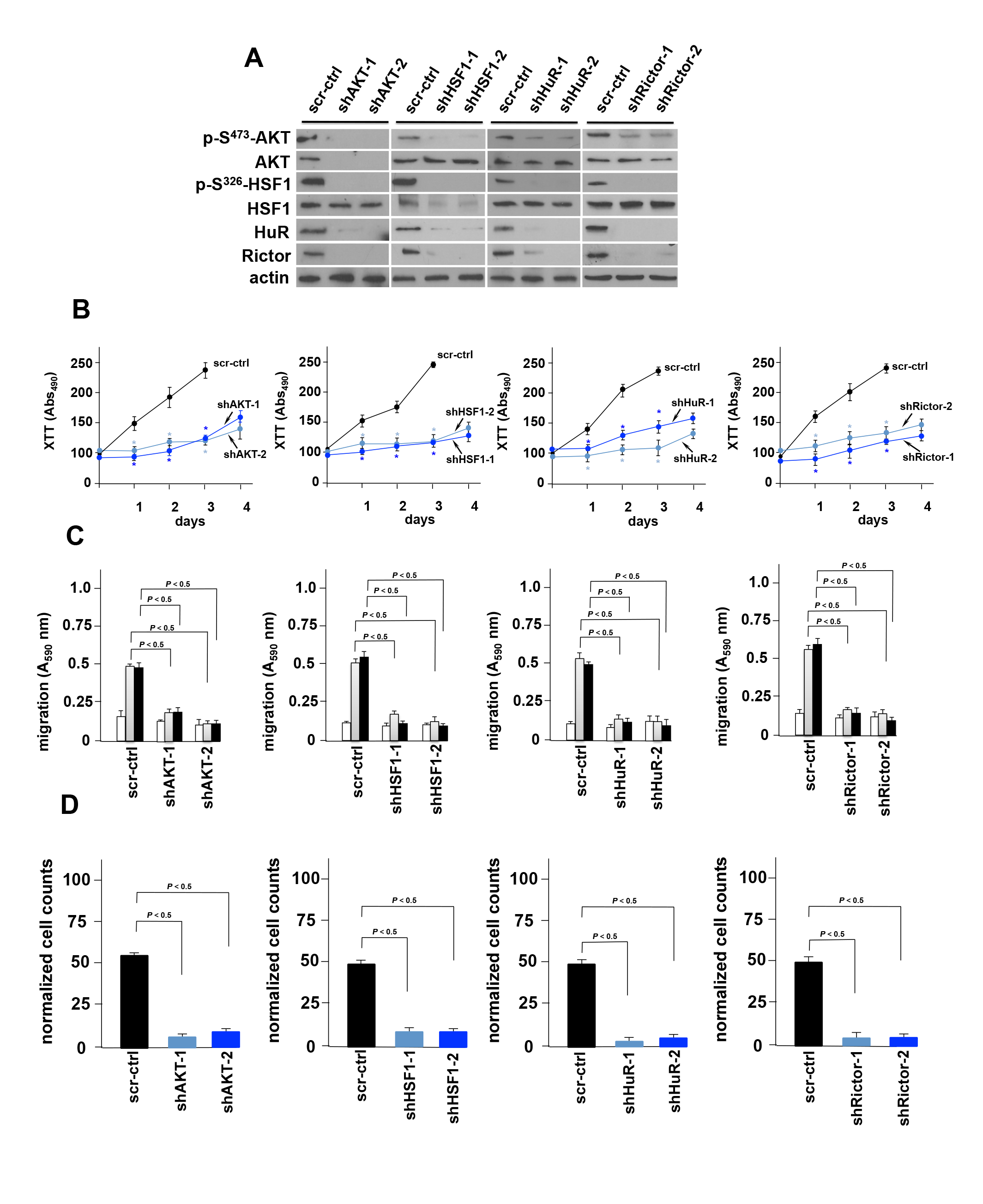
AKT, HSF1 and HuR are required for Rictor expression and their blockade inhibits GBM cell line growth, motility and invasiveness. (**A**) shRNA-mediated knockdown of AKT, HSF1, HuR and Rictor in DBTRG-05MG GBM cells. Cells expressing the indicated shRNA or nontargeting shRNA (negative control, scr; scrambled sequence) were immunoblotted for the indicated proteins. (**B**) Effects of AKT, HSF1, HuR and Rictor knockdown on cell growth in as indicated. Control (scr-ctrl) U87EGFRvIII cells expressed a nontargeting-scrambled shRNA. (*, *P* < 0.05). Mean ± S.D. are shown, n = 3. (**C**) Migration of control (scr-ctrl) or AKT, HSF1, HuR or Rictor shRNA-expressing knockdown clones. Cells were seeded into Boyden chambers and allowed to migrate towards BSA (white bars), vitronectin (grey bars) or fibronectin (black bars). Data represent mean +S.D. of three independent experiments. (**D**) Invasive potential of control or AKT, HSF1, HuR or Rictor shRNA-expressing knockdown cells migrating through matrigel. Data represent mean +S.D. of three independent experiments.

### Constitutive HuR expression prevents loss of Rictor under conditions of AKT or HSF1 knockdown

To gain insight as to whether an mTORC2/phospho-S^473^-AKT/phospho-S^326^-HSF1/HuR/Rictor signaling cascade was operative in GBM, we investigated whether HuR expression driven from a viral vector would prevent Rictor loss under conditions of AKT or HSF1 knockdown. DBTRG-05MG cells were stably transduced with a lentiviral construct which expresses HuR and cells treated with siRNAs targeting AKT, HSF1 or control scrambled non-targeting siRNAs. As shown in figure 5A, viral driven expression of HuR (Lv-HuR) maintained Rictor abundance under conditions of AKT or HSF1 loss. Treatment of cells constitutively expressing viral driven HuR with siRNAs targeting AKT, abolished phospho-S^473^-AKT and total AKT levels, as well as, phospho-S^326^-HSF1 levels, but did not result in reduced Rictor expression (see Figure 5B). Similarly, knockdown of HSF1 in cells expressing viral driven HuR resulted in inhibition of phospho-S^326^-HSF1 and total HSF1, however Rictor expression was maintained. Taken together these data are consistent with a feed-forward loop in which mTORC2 activity leads to signaling through an AKT/HSF1/HuR/Rictor cascade in GBM cells.

**Figure 5.**
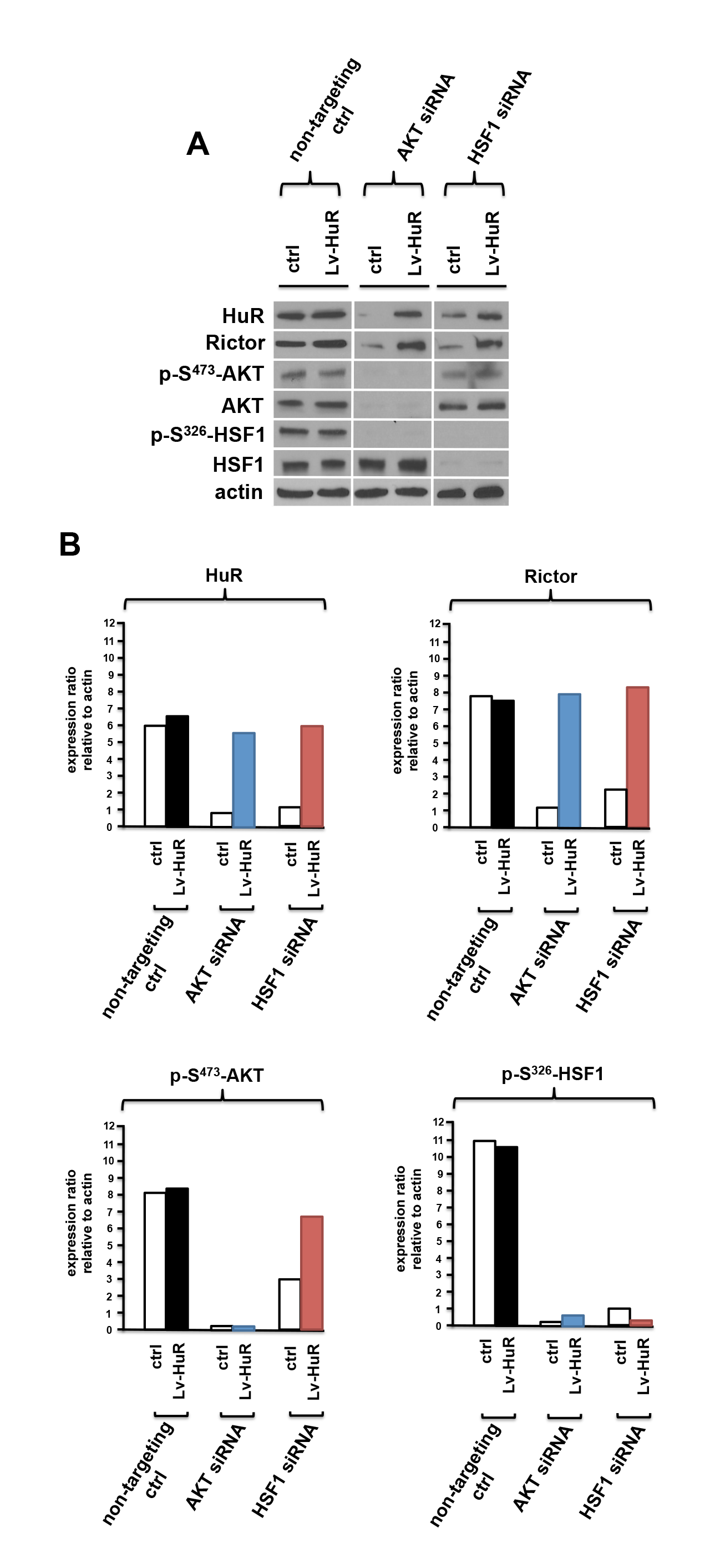
Overexpression of HuR prevents loss of Rictor expression under conditions of AKT or HSF1 loss. (**A**) Viral vector driven expression of HuR prevents inhibition of Rictor expression in DBTRG-05MG cells treated with non-targeting (scr-ctrl), AKT-, or HSF1-targeting siRNAs. Lysates were subsequently immunoblotted for the indicated. (**B**) Quantification of HuR, Rictor, p-S^473^-AKT and p-S^326^-HSF1 protein levels from experiments in (**A**) by densitometry. Experiments were repeated twice with similar results.

### *miR-218*-mediated Rictor repression requires mTORC2/STAT3 signaling

The microRNA *miR-218* negatively regulates Rictor translation in several cancers ^29-31^. *miR-218* is encoded within the intronic sequences of the *SLIT2* and *SLIT3* genes however, it has been shown to mainly originate from the intronic sequences of the *SLIT2* locus in GBM ^32^. Moreover, in U87EGFRvIII-expressing cells it has been shown that STAT3 directly represses *miR-218* transcription by binding to a STAT3 binding site four kilobases downstream of the *miR-218* transcriptional start site (designated the “*miR-218*+4kb” site) ^32^. Recent reports have also suggested a mechanism by which mTORC2-generated signals regulate transcriptional activation of interferon-regulated genes via regulation of STATs ^33^. Thus, we tested whether mTORC2 is involved in the negative regulation of *miR-218* transcription via STAT3. Initially, we confirmed *miR-218* repression in U87EGFRvIII cells relative to parental U87 cells and examined whether *miR-218* was still repressed under conditions of Rictor or mSIN1 loss (Figure 6A). Knockdown of either Rictor or mSIN1 completely blocked the down-regulation of *miR-218* in U87EGFRvIII cells. We further confirmed that Rictor or mSIN1 knockdown did not inhibit *SLIT2* expression (Figure 6B). We subsequently performed a series of ChIP experiments to determine if Rictor or mSIN1 loss affected STAT3 binding to the *miR-218*+4kb binding site. As shown in Figure 6C, in U87EGFRvIII cells transfected with control non-targeting siRNAs, robust STAT3 binding was observed at the *miR-218*+4kb site relative to parental U87 cells. However, STAT3 binding was markedly reduced under conditions of Rictor or mSIN1 loss. These effects were also observed in DNA-pull down assays utilizing beads conjugated with miR-218+4kb DNA sequences. As shown in Figure 6D, nuclear extracts from U87EGFRvIII bound high levels of STAT3, whereas under conditions of Rictor or mSIN1 knockdown STAT3 binding was markedly reduced. These data suggest that mTORC2 is required for STAT3 mediated repression of *miR-218* in GBM cells.

**Figure 6.**
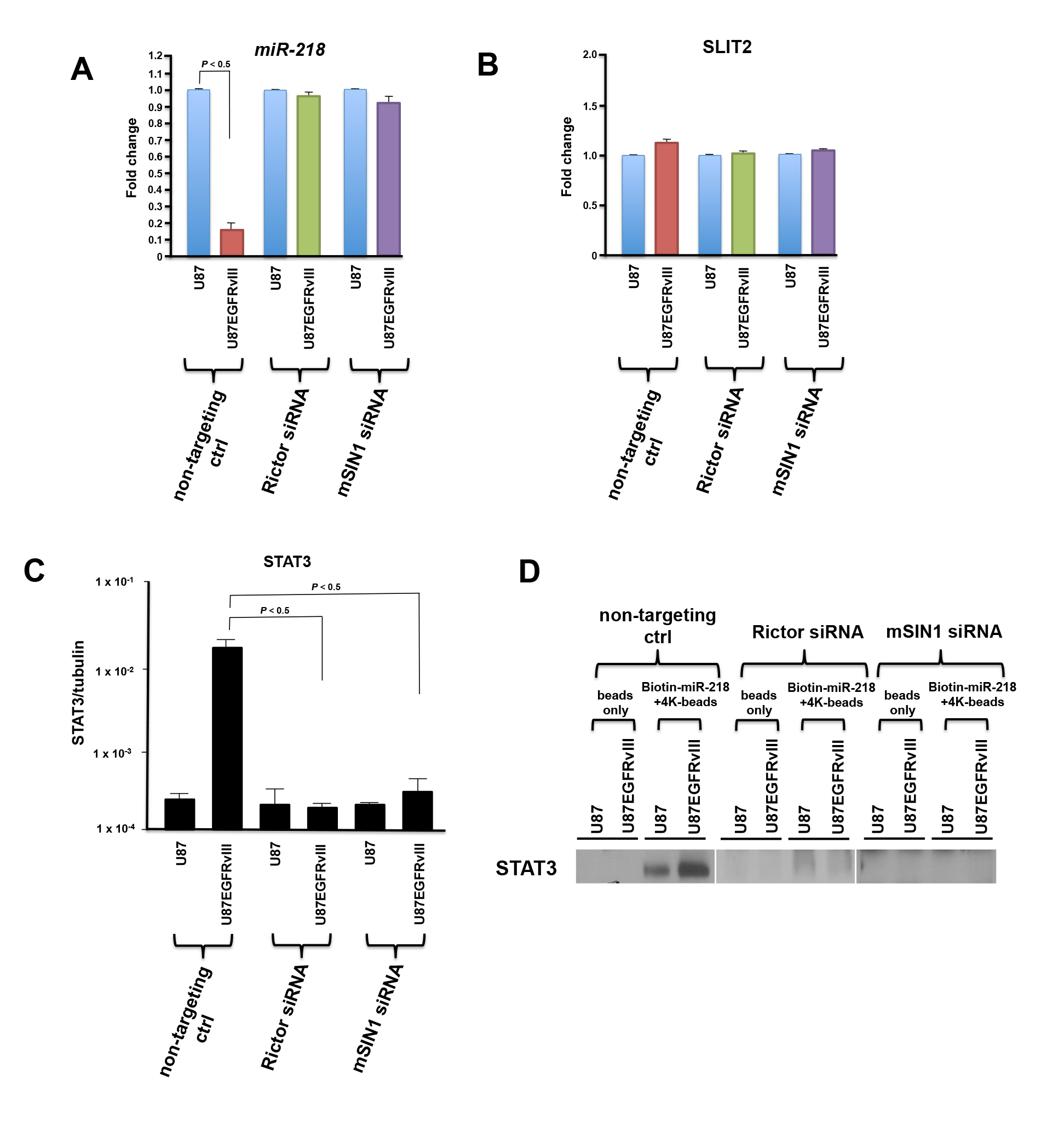
STAT3 dependent *miR-218* repression requires mTORC2 activity in GBM. (**A**) *miR-218* levels in U87 and U87EGFRvIII cells under conditions of Rictor or mSIN1 loss. Cells were treated with either non-targeting (ctrl), Rictor or mSIN1 siRNAs and *miR-218* levels determined. (**B**) Relative levels of SLIT2 upon inhibition of Rictor or mSIN1 expression by siRNA treatments as indicated. (**C**) ChIP analysis for STAT3 binding at the *miR-218*+4kb site in control, Rictor or mSIN-siRNA treated cells as shown. (**D**) *In vitro*-DNA pull-down assay using beads or beads conjugated to *miR-218*+4kb binding site sequences of lysates from the indicated cell lines treated with control (ctrl), Rictor or mSIN1-targeting siRNAs. Experiments were repeated twice with similar results.

### Alterations in mTORC2/AKT/HSF1/HuR signaling affect tumor xenograft growth and Rictor mRNA translation

To determine if shRNA mediated knockdown of AKT, HSF1, or HuR would affect the *in vivo* growth rates of murine xenografts, we subcutaneously implanted DBTRG-05MG cells expressing these shRNAs into SCID mice and monitored tumor growth. As shown in figure 7A, cells expressing non-targeting shRNAs exhibited rapid growth with a latency period of 14 days. Conversely, tumors expressing shRNAs targeting AKT, HSF1, or HuR grew significantly slower and with longer latency periods (*, *P* < 0.05; latency periods; shRNA-AKT, 28 days; shRNA-HSF1, 30 days; shRNA-HuR, 23 days). We also expressed shRNAs targeting Rictor in DBTRG-05MG cells and knockdown resulted in significantly slower growth and longer latency period (shRNA-Rictor, 23 days), confirming our previous results in other GBM cell lines ^16^. Tumors from mice at autopsy were subjected to polysome analyses to determine the relative translational state of the Rictor mRNA. As shown in Figure 7B, in tumors expressing the non-targeting control shRNA 80% of Rictor mRNA was polysome associated and well translated, however in tumors expressing AKT, HSF1 or HuR shRNAs, Rictor mRNA shifted markedly to non-ribosomal/monosomal fractions indicating reduced translational efficiency and consistent with the reduced growth rates of these tumors.

**Figure 7.**
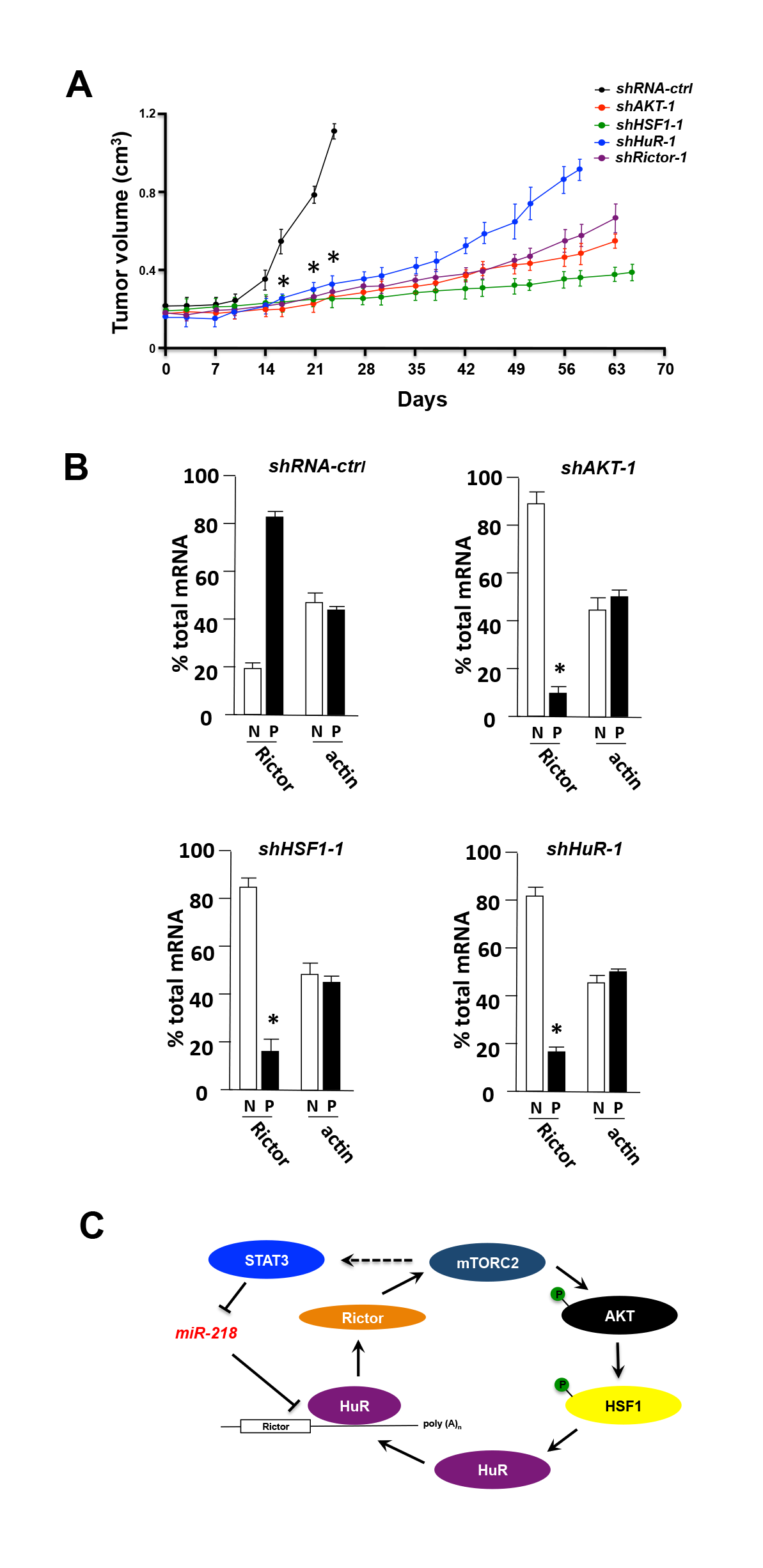
Knockdown of AKT, HSF1 or HuR inhibit GBM growth *in vivo*. (**A**) DBTRG-05MG cells expressing shRNA targeting AKT, HSF1, HuR, Rictor or non-targeting (ctrl) were monitored for tumor growth for up to 70 d. (n = 4-5 per group; *, *P* < 0.05). (**B**) Polysome analyses were performed on cells from tumors harvested at autopsy (as in Figure 1C) for the cells expressing the indicated shRNAs. The distribution of Rictor and actin mRNAs across the gradients was determined and quantified via rt-PCR as before (*, *P* < 0.05). (**C**) Proposed model of feed-forward regulation of Rictor expression and mTORC2 activity via AKT/HSF1/HuR signaling. Additionally, *miR-218* levels are suppressed by elevated mTORC2/STAT3 activity.

### mTORC2/AKT/HSF1/HuR/Rictor signaling in GBM patients

To assess whether these signaling relationships were valid in clinical GBM samples we analyzed an independent set of 34 flash-frozen GBM and 5 normal brain samples. Each tumor sample was confirmed histologically, tumor extracts prepared, and the total relative abundance of phospho-S^473^-AKT, phospho-S^326^-HSF1, HuR and Rictor determined by Western analyses. *miR-218* levels were assessed via quantitative rt-PCR and levels of nuclear STAT3 expression determined via immunohistochemical staining. These data are summarized in table 1. As shown, elevated mTORC2 activity, as determined by immunoblotting for phospho-S^473^-AKT levels, was observed in 22 of 34 tumor samples (65%, *P* < 0.05) consistent with the degree of hyperactivity previously observed ^8, 16^. Phospho-S^326^-HSF1, HuR and Rictor expression was elevated in 74% (25 of 34, *P* < 0.05), 62% (21 of 34, *P* < 0.05) and 74 % (25 of 34, *P* < 0.05) of samples, respectively. Significant direct correlations were observed between samples harboring elevated phospho-S^473^-AKT and increased phospho-S^326^-HSF1, and those containing elevated phospho-S^326^-HSF1 and increased HuR levels (*P* values less then 0.05 for both correlations). A highly significant direct correlation was found between elevated HuR containing samples and those tumors expressing high levels of Rictor (*P* < 0.01). Moreover, elevated mTORC2 activity was directly correlated between increased nuclear STAT3 expression and inversely correlated with elevated *miR-218* levels (*P* < 0.05 and *P* < 0.01, respectively). These data strongly support the feed-forward signaling pathway observed in the GBM cell line experiments and provide evidence of these signaling relationships in patient samples.

**Table 1.** Relative protein levels of phospho-S^473^AKT, phospho-S^326^-HSF1, HuR, Rictor, *miR-218* and nuclear STAT3 in normal and glioblastoma samples.

| Samples | pS473-AKT<br>expression<sup>§</sup> | pS326-HSF1<br>expression<sup>#</sup> | HuR<br>expression<sup>¥</sup> | Rictor<br>expression<sup>¶</sup> | miR-218<br>expression<sup>⌢</sup> | Nuclear STAT3<br>expression |
| --- | --- | --- | --- | --- | --- | --- |
| Normal |  |  |  |  |  |  |
| 1 | 1.1 | 1.6 | 1.9 | 1.5 | 1.1 | -, <20% |
| 2 | 1.7 | 2.1 | 1.3 | 1.1 | 1.5 | -, <20% |
| 3 | 1.2 | 1.3 | 1.4 | 1.4 | 1.2 | -, <20% |
| 4 | 1.6 | 1.4 | 1.8 | 1.9 | 1.6 | -, <20% |
| 5 | 1.5 | 1.8 | 1.7 | 1.2 | 1.3 | -, <20% |
| GBM |  |  |  |  |  |  |
| 1 | 54.3<sup>***</sup> | 26.3<sup>++</sup> | 37.5<sup>***</sup> | 65.9<sup>«++</sup> | 13.5<sup>++</sup> | ++, 20-50% |
| 2 | 15.7<sup>+</sup> | 19.7<sup>+</sup> | 8.1 | 10.2<sup>«+</sup> | 2.5 | -, <20% |
| 3 | 26.9<sup>***</sup> | 17.3<sup>++</sup> | -<sup>‡</sup> | 49.4<sup>«++</sup> | 6.3 | +, 20-50% |
| 4 | 36.2<sup>***</sup> | 43.8<sup>++</sup> | 9.62 | 54.1<sup>«++</sup> | 2.1 | ++, >50% |
| 5 | 48.4<sup>***</sup> | 34.3<sup>++</sup> | 42.7<sup>***</sup> | 26.8<sup>«++</sup> | 4.2 | ++, >50% |
| 6 | 3.5 | 0.7 | 2.7 | 1.8 | 16.8<sup>++</sup> | -, <20% |
| 7 | 26.9<sup>***</sup> | 16.4<sup>***</sup> | 53.1<sup>***</sup> | 29.3<sup>«++</sup> | 1.7 | +, >50% |
| 8 | 5.5 | 2.4 | 3.9 | 6.9 | 19.2<sup>++</sup> | -, <20% |
| 9 | 32.4<sup>***</sup> | 45.9<sup>++</sup> | 57.3<sup>***</sup> | 48.2<sup>«++</sup> | 2.9 | ++, 20-50% |
| 10 | 23.6<sup>***</sup> | 29.7<sup>++</sup> | 42.8<sup>***</sup> | 37.4<sup>«++</sup> | 5.6 | ++, 20-50% |
| 11 | 17.6<sup>+</sup> | 11.3<sup>+</sup> | 13.1<sup>+</sup> | 9.7<sup>«+</sup> | 3.7 | +, 20-50% |
| 12 | 39.1<sup>***</sup> | 56.2<sup>***</sup> | 64.5<sup>***</sup> | 25.8<sup>«++</sup> | 1.3 | ++, >50% |
| 13 | 4.3 | 13.5<sup>+</sup> | 8.2 | 3.4 | 10.5<sup>++</sup> | -, <20% |
| 14 | 32.7<sup>***</sup> | 26.9<sup>++</sup> | 29.2<sup>***</sup> | 41.9<sup>«++</sup> | 2.7 | +, 20-50% |
| 15 | 56.8<sup>***</sup> | 32.8<sup>++</sup> | 46.9<sup>***</sup> | 41.0<sup>«++</sup> | 1.9 | ++, >50% |
| 16 | 3.8 | -<sup>‡</sup> | 4.2 | 2.1 | - | -, <20% |
| 17 | 7.8 | 10.3<sup>+</sup> | 11.3<sup>+</sup> | 7.5 | 16.4<sup>++</sup> | -, <20% |
| 18 | 42.7<sup>***</sup> | 38.6<sup>++</sup> | 52.6<sup>***</sup> | 31.1<sup>«++</sup> | 4.6 | ++, >50% |
| 19 | 27.9<sup>***</sup> | 29.4<sup>++</sup> | 24.8<sup>***</sup> | 30.4<sup>«++</sup> | 3.2 | ++, >50% |
| 20 | 43.6<sup>***</sup> | 48.1<sup>++</sup> | 39.7<sup>***</sup> | 37.9<sup>«++</sup> | 5.8 | +, >50% |
| 21 | 6.7 | 5.3 | 4.6 | 3.6 | 12.4<sup>++</sup> | -, <20% |
| 22 | 46.1<sup>***</sup> | 26.7<sup>++</sup> | 38.6<sup>***</sup> | 42.8<sup>«++</sup> | 2.2 | ++, 20-50% |
| 23 | 7.2 | 5.4 | -<sup>‡</sup> | 2.4 | 9.7 | -, <20% |
| 24 | 78.4<sup>***</sup> | 69.4<sup>++</sup> | 52.8<sup>***</sup> | 34.8<sup>«++</sup> | 2.4 | ≈ |
| 25 | 53.2<sup>***</sup> | 66.8<sup>++</sup> | 29.8<sup>***</sup> | 53.1<sup>«++</sup> | 4.8 | ++, 20-50% |
| 26 | 69.2<sup>***</sup> | 13.4<sup>++</sup> | 42.5<sup>***</sup> | 16.4<sup>«+</sup> | 6.1 | ++, >50% |
| 27 | 7.2 | 5.2 | 3.7 | 2.6 | 17.3<sup>++</sup> | -, <20% |
| 28 | 76.3<sup>***</sup> | 84.3<sup>++</sup> | 63.8<sup>***</sup> | 58.4<sup>«++</sup> | 2.1 | ++, 20-50% |
| 29 | 7.0 | 2.4 | 6.4 | 5.8 | 9.9 | -, <20% |
| 30 | 16.2<sup>+</sup> | 14.7<sup>+</sup> | 9.1 | 12.5<sup>«+</sup> | 18.4<sup>++</sup> | +, <20% |
| 31 | 43.8<sup>***</sup> | 38.6<sup>++</sup> | 5.5 | 49.2<sup>«++</sup> | 1.3 | ++, 20-50% |
| 32 | 62.5<sup>***</sup> | 39.4<sup>***</sup> | 46.4<sup>***</sup> | 58.6<sup>***</sup> | 9.5 | ++, 20-50% |
| 33 | 28.4<sup>***</sup> | 51.3<sup>***</sup> | 33.5<sup>***</sup> | 32.9<sup>«++</sup> | 5.4 | ++, >50% |
| 34 | 48.1<sup>***</sup> | 50.7<sup>++</sup> | 43.7<sup>***</sup> | 29.6<sup>«++</sup> | 4.9 | ++, 20-50% |

Note: Five normal brain and 34 independent quick-frozen GBM samples were assessed for HuR, Rictor, phosphorylated HSF1 and phosphorylated AKT levels by Western analyses as described in Experimental Procedures and quantified by densitometry. 22 of 34 tumor samples (65%) had markedly higher mTORC2

## DISCUSSION

Our current understanding suggests that mTORC2 can be regulated by RTKs, ribosomes, TSC1-TSC2, Rac1, and the expression levels of Rictor ^2, 4^ Our data suggest that mTORC2 is also subject to autoregulation via feed-forward signaling through an AKT/HSF1/HuR/Rictor cascade in GBM (see Figure 7C). Additionally, our data suggests that mTORC2 activity regulates STAT3 binding to the *miR-218*+4kb binding site and represses *miR-218* transcription, thereby negating its suppressive effects targeting the Rictor mRNA.

Although Rictor is necessary for the stability and activity of mTORC2 and reports have described the role of phosphorylation and acetylation events in these processes, little is known regarding the control of Rictor expression via post-transcriptional control mechanisms ^4, 34^ The 3′ UTR of Rictor is relatively long suggesting that it contains post-transcriptional regulatory sequences. Furthermore, the 3′ UTR contains several segments of AU-or U rich *cis*-regulatory sequences implicated in mRNA turnover and translational control ^35, 36^. Indeed, our analysis identified several HuR binding sites (see figure 2A) and additional regulatory domains are likely to be present in such a large 3′ UTR, possibly controlling mRNA stability under particular conditions. Our data supporting the ability of the Rictor transcript to be subject to translational control is also reinforced by recent data from Yasuda and colleagues, who investigated the role of Mdm20 in actin remodeling via Rictor-mediated mTORC2 activity ^20^. Mdm20 was suggested to regulate the expression of Rictor at the level of *de novo* protein synthesis.

Our study implicates HSF1/HuR signaling in the regulation of Rictor expression and mTORC2 activity. Recent data support an HSF1/HuR cascade in the regulation of β-catenin levels in breast cancers ^25^. Gabai *et al*, also observed that HSF1 controls the expression of HIF-1α via effects on HuR ^37^. HSF1’s affects on HuR were determined to be at the level of transcription. We identified tandem non-canonical HSEs within the promoter of HuR and showed that these sites were capable of association with activated HSF1 and promoted HuR expression as determined by increased *Pol* II occupancy (see Figure 3). These sequences were also identified as HSF1 binding sites in a high-resolution survey of HSF1 genome occupancy in breast cancer cells ^28^. These data support the notion that HuR is a direct target of activated HSF1 in GBM.

We observed that mTORC2 activity was required for STAT3 dependent *miR-218* repression in GBM ^32^. STAT3 predominantly functions as a transcriptional activator, however several studies have demonstrated that STAT3 can also act as a repressor ^38, 39^. Loss of Rictor or mSIN1 was sufficient to block *miR-218* repression, restoring *miR-218* to control levels (see Figure 6A). We attempted to ascertain whether in U87EGFRvIII cells, increased phosphorylated STAT3 (either phospho-Y^705^ or phospho-S^727^) could be correlated with elevated phospho-Y^705^/S^727^-STAT3 binding to the *miR-218*+4kb binding site, however neither phospho-Y^705^ or phospho-S^727^ STAT3 were found bound to the *miR-218*+4kb site in U87EGFRvIII cells as determined via DNA pull-down assays (not shown). These observations suggest that mTORC2 may phosphorylate STAT3 at a distinct phospho-site(s) resulting in binding of STAT3 to the *miR-218*+4kb site and repressing *miR-218* transcription. Alternatively, via an indirect mechanism, mTORC2 may recruit ancillary factors which together with STAT3 mediate *miR-218* repression. Along these lines, Mathew *et al*., identified BCLAF1 as displaying increased recruitment to the *miR-218*+4kb binding region ^32^. BCLAF1 is a transcriptional repressor and has been shown to interact with antiapoptotic components of the BCL2 family ^40^. Additionally, BCLAF1 has also been shown to interact with BRCA1 which can lead to the formation of a BRCA1-mRNA splicing complex in response to DNA damage ^41^. Thus, BCLAF1 mediated recruitment of the BRCA1-splicing machinery favors pre-mRNA splicing of target genes. Mathew *et al*., suggested that an intronic region spanning the *miR-218* locus may be preferentially spliced out via a BCLAF1-dependent co-transcriptional splicing event ^32^. It is conceivable that such preferential splicing may be regulated by mTORC2 activity. Further studies will be required to elucidate these mechanistic details.

All of the components of the signaling cascade we have delineated have established roles in GBM growth and migration. Rictor/mTORC2 have important roles regulating these functions and additionally regulate the metabolic reprograming of these tumors ^2, 16^. AKT and HSF1 have been shown to mediate glioma growth and survival ^42, 43^, while HuR appears to be overexpressed in GBM and its dysregulation may result in the enhanced translation of mRNAs promoting growth, motility and drug resistance ^44^ Feed-forward signaling loops are well represented in cell signaling networks and provide a positive mechanism for reinforcing and stimulating network activity. Many of the downstream effectors of the feed-forward loop components described here have overlapping targets and the coordinated activation of this loop may serve to stimulate the expression of common targets at multiple levels of gene regulation. These results contribute to a better understanding of the signaling mechanisms regulating Rictor expression and mTORC2 activity in GBM.

## MATERIALS & METHODS

### Cell Lines, GBM samples, Transfections and Viral Transductions

All GBM cell lines were obtained from ATCC except U87EGFRvIII, which was kindly provided by Dr. Paul Mischel (LICR-UCSD). Flash-frozen normal brain and glioblastoma samples were obtained from the Cooperative Human Tissue Network under an approved Institutional Review Board protocol. Each sample was histopathologically reviewed and those containing greater than 95% tumor were utilized. Samples were homogenized in RIPA buffer using a Polytron homogenizer (Fisher, Pittsburgh, PA) to generate extracts for protein and RNA analysis. Sections of paraffin-embedded tumors on slides were processed for immunohistochemistry as previously described ^16^. DNA transfections were performed using Effectene transfection reagent according to the manufacturer (Qiagen). For siRNA knockdowns, DBTRG-05MG, U87 or U87EGFRvIII cells were transfected with 10 nmol/L siRNA pools targeting AKT1, HSF1, HuR, Rictor, mSIN1 or a non-targeting scrambled control sequence. ON-TARGETplusSMARTpool siRNAs were obtained from GE Dharmacon and transfected using Lipofectamine RNAimax (Life Technologies). Lentiviral shRNA production and infection was performed as previously described ^16^.

### Constructs and Reagents

The luciferase-human Rictor 3′ UTR constructs were generated by insertion of the full-length human Rictor 3′ UTR into the *Xba*I site of pGL3-promoter (Promega). Mutagenesis was performed using the QuikChange Lightning Multi Site-Directed Mutagenesis Kit (Agilent Technologies) with the appropriate mutagenic primers according to the manufacturer. TRC pLKO.1 library constructs expressing multiple shRNA-targeting AKT1, HSF1, HuR, Rictor or non-targeting controls were obtained from GE Dharmacon. The constructs utilized had the following TRC designations: shAKT-1, TRCN0000010162; shAKT-2, TRCN0000010163; shHSF1-1, TRCN0000007480; shHSF1-2, TRCN0000007481; shHuR-1, TRCN0000017273; shHuR-2, TRCN0000017274; shRictor-1, TRCN0000074288; shRictor-2, TRCN0000074289. The HuR overexpression construct contained the full-length human HuR ORF cloned into pLenti-C-Myc-DDK-P2A-Puro (Lv-HuR) and was from Origene. Antibodies to the following proteins were used: phospho-S^473^-AKT (#9271, CST), AKT (#9272, CST), Rictor (#A300-459A, Bethyl Laboratories), actin (#ab3280, Abcam), phospho-Y^1068^-EGFR (#2234, CST), *β*-tubulin (#2146, CST), phospho-S^326^-HSF1 (ADI-SPA-902-D, Enzo Life Sciences), HSF1 (ADI-SPA-901-D, Enzo Life Sciences), HuR (07-468, EMD Millipore), RNA Pol II (#39097, Active Motif) and STAT3 (#4904, CST). AG1478 was obtained from Selleckchem. EGF was from Life Technologies and all other reagents were from Sigma.

### Polysome Analysis

Separation of polysomes was performed as previously described ^45^. Briefly, cell extracts were prepared and layered onto 15% to 50% sucrose gradients and spun at 38,000 rpm for 2 h at 4°C in a SW40 rotor (Beckman Instruments). Gradients were fractionated using a gradient fractionator system (Brandel Instruments) using a flow rate of 3 mL/min. The polysome profile of the gradients was monitored via UV absorbance at 260 nm. RNA was isolated and pooled into nonribosomal/monosomal and polysomal fractions. RNAs (100 ng) were subsequently used in quantitative reverse transcriptase-PCR analyses.

### Immunoblotting and Quantitative real time PCR

Western blotting was performed as previously described ^46^. miRNeasy mini kit (Qiagen) was used to isolate total RNA and the RNA was reverse transcribed into cDNA using the High Capacity RNA-to-cDNA kit (ABI). Analysis of *miR-218* expression was performed using the TaqMan MicroRNA Reverse Transcription kit (ABI ThermoFisher) according to the manufacturers instructions. Taqman primers (from Applied Biosystems) were used to measure the levels of all transcripts and analyses were performed on an ABI7900HT system (ABI ThermoFisher).

### Chromatin immunoprecipitations, RNA and DNA *in vitro* pull-down assays

Chromatin immunoprecipitation (ChIP) assays were performed as previously described ^47^. For RNA-pull down assays ^48^, extracts were prepared and biotinylated RNA oligonucleotides containing HuR binding site motif(s) were added. The protein and biotinylated RNA complexes were recovered and the complexes were washed five times and resolved by gel electrophoresis. *In vitro* DNA-pull down assays were performed as described ^47^ using a double-stranded oligonucleotide containing the human HuR noncanonical HSE or the *miR-218*+4kb binding site attached to streptavidin-Sepharose beads via a 5’ biotinylated plus strand according to the manufacture’s recommendation (Invitrogen). Beads with bound proteins were analyzed by SDS-PAGE followed by immunoblotting.

### Cell proliferation and migration assays

Cells growth was determined via XTT assays (Roche). Cell migration assays were performed using modified Boyden chambers (Chemicon) as previously described ^49^. For invasion assays through Matrigel, 2 × 10^4^ cells were placed into the top well of Boyden chambers containing growth factor-reduced Matrigel extracellular basement membrane over a polyethylene terephthalate membrane (8-mm pores; BD Biosciences). Following 24 h culture, Matrigel was removed and invaded cells were fixed and stained. Cells adhering to the bottom of the membrane were counted.

### Xenograft studies

All animal experiments were performed under an approved Institutional Animal Care and Use Committee protocol and conformed to the guidelines established by the Association for the Assessment and Accreditation of Laboratory Animal Care. Xenografts of shRNA-expressing cell lines were injected s.c. into the flanks of 4-6 week old female C.B.-17-scid (Taconic) mice as previously described ^16^. Tumors were measured every 3-4 days and tumor volumes calculated using the formula length × width × height × 0.5236. Tumors were harvested at autopsy for polysome analyses.

### Statistical analysis

Statistical analyses were performed using unpaired Student’s *t* tests and ANOVA models using Systat 13 (Systat Software, Chicago, IL). *P* values of less the 0.05 were considered significant.

## ACKNOWLEDGMENTS

We thank Drs. Jacob Fleischmann, Norimoto Yanagawa, Sanjai Sharma and Paul Mischel for cell lines and reagents. We also thank Dr. Alan Lichtenstein for comments on the manuscript and Jheralyn Martin and Janine Masri for technical assistance. We are also grateful to Dr. Denson Fujikawa for assistance with tissue processing, embedding and sectioning. This work was supported, in whole or in part, by VA MERIT I01BX002665 and NIH R01CA109312 grants.

## Author Contributions

Conceived and designed the experiments: BH, RNN, JG. Performed the experiments: BH, ABS, RSF, KAL, TB. Analyzed the data: BH, RNN, JG. Wrote the paper: BH, RNN, JG.

